# Symmetric Intralimb Transfer of Skilled Isometric Force Production

**DOI:** 10.1101/491415

**Authors:** Vikram A Rajan, Robert M. Hardwick, Pablo A Celnik

## Abstract

Motor control theories propose that the same motor plans can be employed by different effectors. Skills learned with one effector can therefore ‘transfer’ to others, which has potential applications in clinical situations. However, evidence from visuomotor adaptation suggests this effect is asymmetric; learning can be generalized from proximal-to-distal effectors (e.g. arm to hand), but not from distal-to-proximal effectors (e.g. hand to arm). We propose that skill learning may not be subject to this asymmetry, as it relies on multiple learning processes beyond error detection and correction. Participants learned a skill task involving the production of isometric forces. We assessed their ability to perform the task with the hand and arm. One group trained to perform the task using only their hand, while a second trained using only their arm. In a final assessment, we found that participants who trained with either effector improved their skill in performing the task with both their hand and arm. There was no change in a control group that did not train between assessments, indicating that gains were related to the training, not the multiple assessments. These results indicate that in contrast to visuomotor adaptation, motor skills can generalize from both proximal-to-distal and distal-to-proximal effectors. We propose this is due to differences in the processes underlying skill acquisition in comparison to visuomotor adaptation.

**New and Noteworthy:** Prior research indicates that motor learning transfers from proximal-to-distal effectors, but not vice-versa. However, this work focused on adapting existing behavior; we questioned whether different results would occur when learning new motor skills. We found that the benefits of training on a skill task with either the hand or arm transferred across both effectors. This highlights important differences between adaptation and skill learning, and may allow therapeutic benefits for patients with impairments in specific effectors.

## Introduction

Understanding how the central nervous system represents learned actions is a central question in motor control research. The theory of generalized motor programs (Schmidt 1975) proposes that learned actions can be generalized to suit the environmental requirements of the task. This suggests that actions, rather than being represented at the level of specific effectors (e.g. arm, hand, etc.) instead share a more abstract control policy that can be modified to account different contexts. A classic example of such generalization is that individuals show similar patterns when writing with their dominant hand compared to when writing with other effectors such as their non-dominant hand or their feet (Schmidt and Lee 2005).

Given that action representations are effector-independent, it should be possible to acquire a control policy for a task with one effector (e.g. the arm), then generalize this policy to improve performance of the same task with an untrained effector (e.g. the hand). This feature might create therapeutic opportunities; a general movement policy developed using one effector could be used to enhance control of an impaired effector(Raghavan et al. 2010). The reverse relationship - whereby training on a task with the impaired effector leads to small but significant improvements in performance with the unimpaired effector - has recently been shown (Kitago et al. 2015). There is, however, evidence that such transfer is asymmetric. Previous studies using visuomotor adaptation tasks indicate that actions can transfer from proximal-to-distal effectors, such as from the arm to the hand, but not from distal-to-proximal effectors (Putterman et al. 1969; Hay and Brouchon 1972; Krakauer et al. 2006). This asymmetry has previously been attributed to the different biological constraints and history of use of these effectors (Krakauer et al. 2006); all movements of the arm affect the position of the hand in space, but not all hand movements similarly affect the arm. Thus, learning relevant actions with the arm is more likely to be transferable to the hand, while learning with the hand is more likely to be effector specific.

Notably, there is limited evidence of asymmetric transfer beyond visuomotor adaptation paradigms. The learning process underlying adaptation requires the simple adjustment of executed actions with the goal of minimizing perturbation-induced error; in contrast, skill learning tasks require the development and refinement of a control policy for actions themselves (Krakauer 2009). As such, skill learning paradigms may allow for symmetric transfer due to the generation of a novel, centrally stored control policy. Here we tested this theory by training participants to perform a task with either the hand or arm, assessing performance with both effectors at different points in the learning process.

## Methods

A total of 30 healthy volunteers (21 female, mean age 23 years) completed the study. Participants were split evenly across three groups with no significant differences in age or sex (one way ANOVA, p>0.9). All participants were right handed as determined by the Edinburg Handedness Inventory, and had no history of neurological conditions. The procedures of the study were approved by the Johns Hopkins Institutional Review Board, and all participants gave informed written consent.

Participants sat in a KINARM robotic exoskeleton (BKIN Technologies, Kingston, Ontario, Canada) positioned in front of a horizontal projector and mirror system. The right arm was kept horizontal to the ground, with the shoulder and elbow fixed at a 45 degree and 90 degree angle, respectively (Figure 1A).

**Figure 1:**
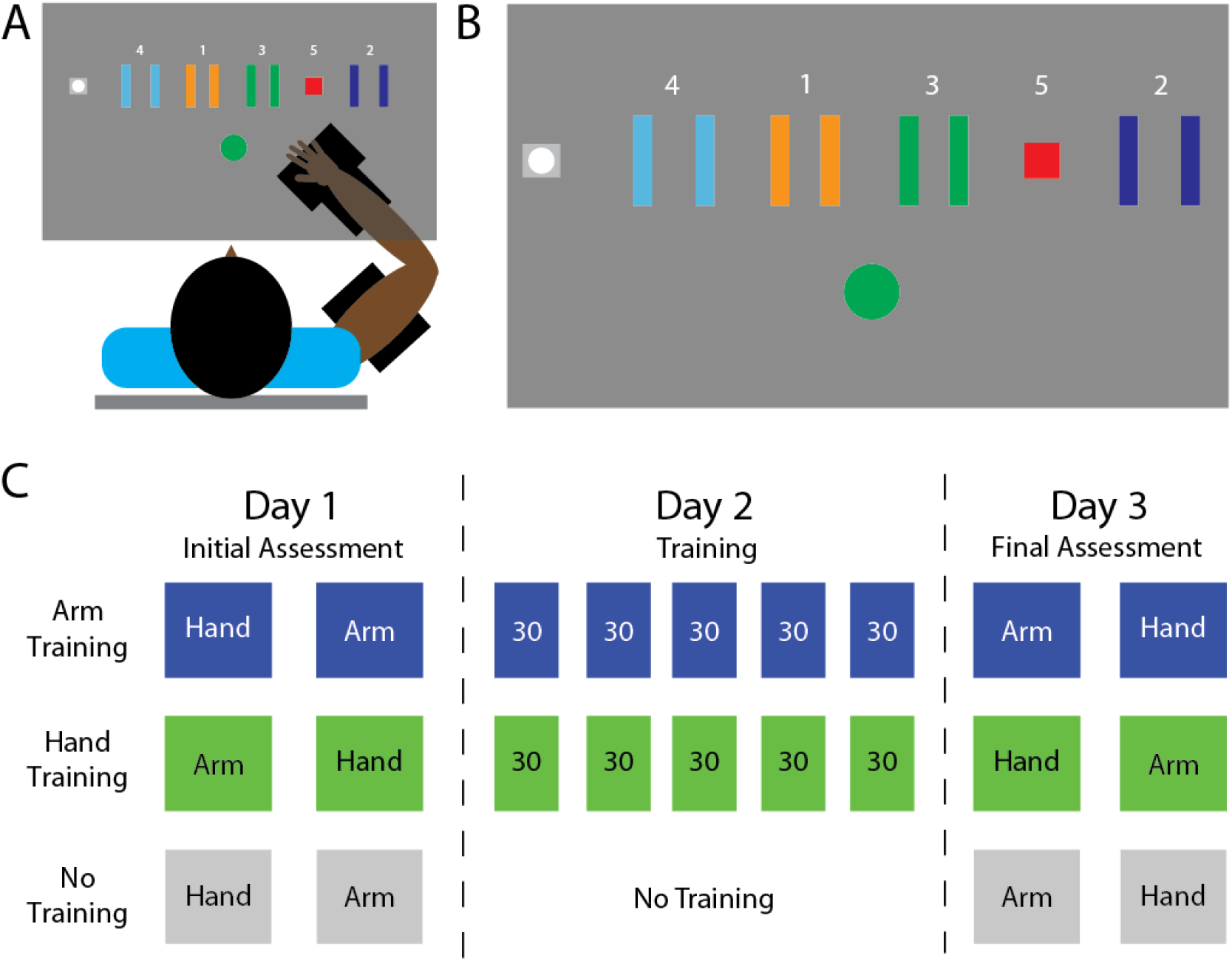
Experimental setup and conditions. A) Participants sat with their arm supported by a robotic exoskeleton. They produced forces with either the hand or arm to control the position of an on screen cursor, attempting to navigate the cursor through a sequence of targets. B) Illustration of the visual display and targets. C) Illustration of the assessment and training schedule for each group of participants during the study.

### Study Design

Participants completed a modified version of the sequential visual isometric pinch task (SVIPT) (Hardwick et al. 2017). They controlled an on-screen cursor by performing isometric contractions, either by pinching a force transducer between the thumb and index finger, or by using their elbow flexor muscles. Of note, the arm and hand were fixed at the same position/angles when performing the task with either effector.

Applying forces moved the cursor horizontally to the right, while relaxing moved the cursor back towards the ‘Home’ position. Numbered targets were arranged horizontally to the right of the Home position (Figure 1B), and participants were instructed to move the cursor through the sequence: “Home-One-Home-Two-Home-Three-Home-Four-Home-Five”. The arrangement of the targets was fixed throughout the study. Force transducer input was passed through a 20 Hz low-pass Butterworth filter to dampen high frequency noise.

As in previous studies using this paradigm, the difficulty of the task was increased by introducing a logarithmic transformation between the applied force and the displacement of the cursor. The required forces were normalized such that a contraction of 30% of the maximum for the effector being used would displace the cursor by 30cm on the screen. This normalization procedure made the task comparable between the two effectors; at 30% of MVC neither arm flexor muscles nor intrinsic hand muscles recruit all available motor units, and are therefore activated in a similar manner (Kukulka and Clamann 1981).

### Skill Assessments

Changes in performance were examined in ‘skill assessments’, which empirically quantified the speed-accuracy trade-off. This allows comparison of performance across groups by controlling for the potentially confounding factors of differences in speed (Reis et al. 2009; Hardwick et al. 2017). Participants completed an initial skill assessment for both the hand and arm on day one of the study and a final skill assessment for both effectors on day three (Figure 1C). Participants were required to complete the sequence at a pace set by an auditory metronome. The tempos used were 24, 30, 38, 45, 60, 80, 100, 110, and 120 beats per minute, translating to approximate trial durations of 12.5,10.0, 7.9, 6.7, 5.0, 3.8, 2.7, and 2.5 seconds respectively. Participants completed 10 trials at each of the 9 tempos in each assessment. The order in which each tempo was attempted was randomized to prevent order effects.

### Training

On day two of the study, the training groups practiced performing the task with either their hand or their arm. As in previous studies using this paradigm, and in contrast to the skill assessments, participants were able to perform the task at a self-selected speed (Reis et al. 2009, 2009; Cantarero et al. 2013a, 2013b; Saucedo Marquez et al. 2013; Statton et al. 2015; Wymbs et al. 2016; Hardwick et al. 2017; Spampinato and Celnik 2017, 2018). Participants were encouraged to perform the task as quickly and as accurately as they could, and to attempt to improve their performance of the task throughout the session. Participants completed 5 blocks of training, each comprising 30 trials.

### No Training Control Group

A third group of participants acted as a control for the possibility that completing skill assessments themselves led to an increase in participant skill. This group completed an initial skill assessment on day one of the study, but did not train before their final assessment on day three. Thus any changes in their performance could only be attributed to acquiring skill during the assessments themselves.

### Data Analysis

For skill assessments, success rates were computed as the proportion of trials within a block in which the participant hit all five targets. The mean average success rate was calculated across the nine movement tempos comprising each skill assessment. Performance in skill assessments was then examined using a mixed-design ANOVA with within-subject factors of *effector* (hand, arm) and *assessment* (initial, final), and a between-subjects factor *group* (hand training, arm training, no training). Based on our hypothesis that skill acquisition would transfer across both effectors (but would not occur for participants in the control group), we conducted planned t-tests, comparing performance on the initial and final skill assessments for each group, split according to the effector used (Bonferroni corrected to account for the 6 comparisons being made, adjusted alpha p<0.0083). Differences in the change in performance from the initial to the final skill assessment were examined with separate ANOVAs to assess hand skill and arm skill for the three groups, with significant difference between groups being examined using t-tests (bonferroni corrected to account for the 3 comparisons being made, adjusted alpha p<0.017).

For training data, online changes in performance within group were examined by conducting paired-samples t-tests on the first and final block of practice for each group. Separate analyses were conducted for success rates and trial durations.

Results from the skill assessments are presented in Figure 2. Participants performed the task with greater accuracy with their hand than their arm (main effect of effector, F_1,27_=32.32, p<0.001). There were no differences between group performance in the initial hand or arm skill assessment (one wayANOVÂs, both F<0.5, p>0.6).

**Figure 2:**
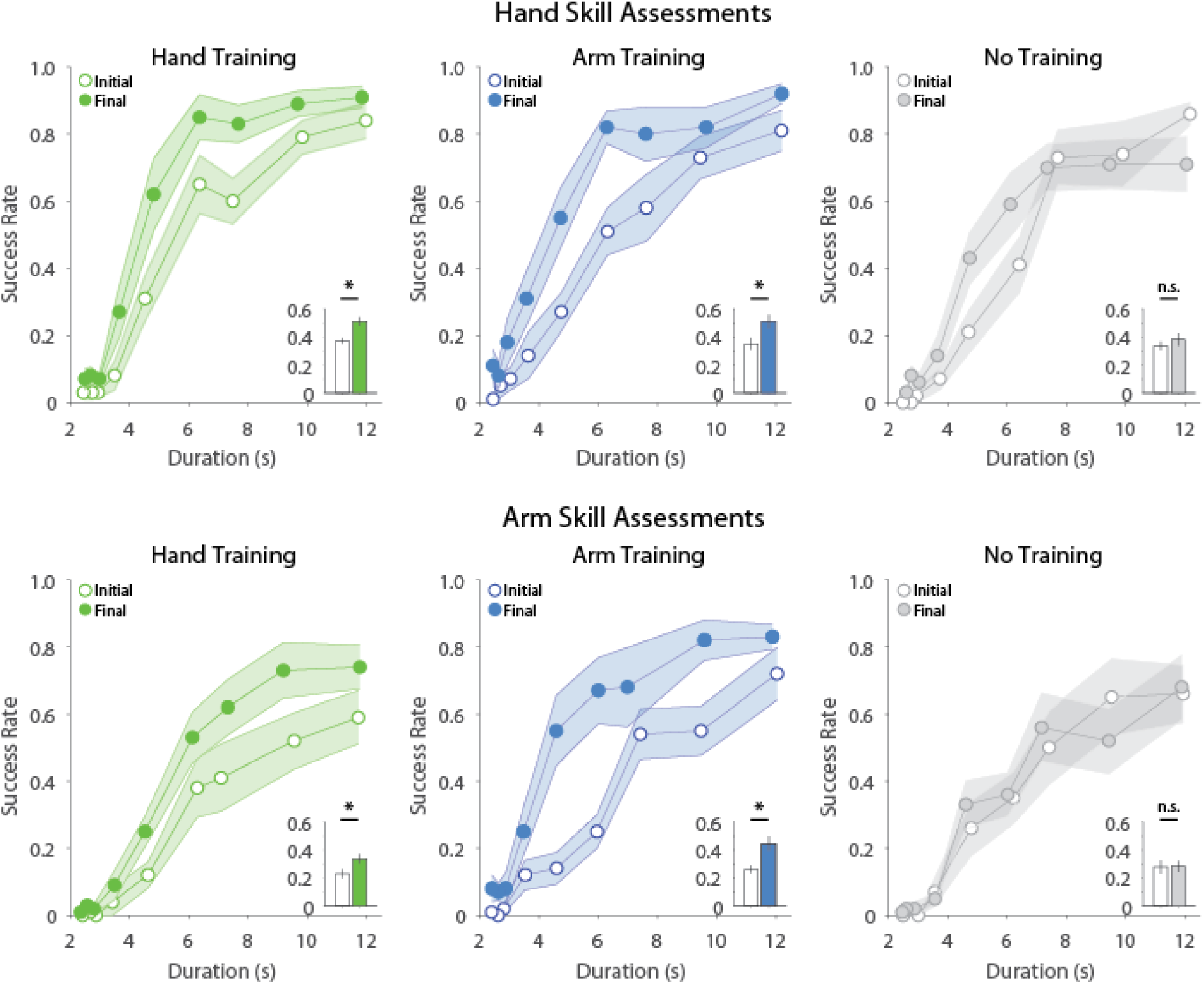
Results of the initial and final skill assessments. Clear and filled circles/bars present data for initial and final skill assessments, respectively. Lines present the speed-accuracy trade-off data for each of the nine movement speeds tested in each assessment. Insets show summary data (average across the nine time points) and statistics. All error bars present ±1SEM. * Indicates p<0.0083

Participants in groups who trained to perform the task with either effector performed better in the final skill assessments compared to the control group (significant assessment x group interaction, F2,27= 13.49, p<0.001, all other interactions n.s.). Participants in groups that trained were able to improve their accuracy with both their trained and untrained effectors by the final skill assessment (paired samples t-tests on initial vs. final skill assessment were all bellow bonferonni corrected critical alpha; p<0.0083). Participants in the no training group did not improve their performance with either effector (paired samples t-tests on initial vs. final skill for hand and arm skill assessments, both p>0.0083).

Additional analyses examined whether the change in performance differed between groups. For hand skill assessments, both training groups had significantly greater improvements in performance than the no training control group (ANOVA on change in hand skill, significant difference between groups, F_2,29_=5.618, p<0.01; bonferroni corrected t-test comparing hand training vs control groups, t_18_=2.69, p<0.017, and arm training vs control groups, t_18_=3.32, p<0.01). However, the amount by which performance improved did not differ between the group that trained with the hand compared to the group that trained with the arm (bonferroni corrected t-test, t_18_=0.54, p>0.017). Similarly, in the final arm skill assessment, both training groups improved their performance significantly more than the no-training group (ANOVA on change in arm skill, significant difference between groups, F_2,29_=10.782, p<0.001; bonferroni corrected t-tests for hand training vs control group, t_18_=3.23, p<0.01, and arm training vs control group, t_18_=4.40, p<0.001). Again, there was no difference between the amount of improvement for the groups that trained with the hand or the arm. (bonferroni corrected t-test, t_18_=1.84, p>0.017).

### Training Data

During the training session on day two, both the hand and arm training groups improved the speed at which they completed the task (paired samples t-tests on initial vs final block of training, hand training group: t_9_=3.55, p<0.01, arm training group, t_9_=3.81, p<0.01). Neither training group changed the accuracy with which they completed the task (paired samples t-test on initial vs final block of training, hand training group: t_9_=0.51, p>0.05, and arm training group, t_9_=0.44, p>0.05).

**Figure 3:**
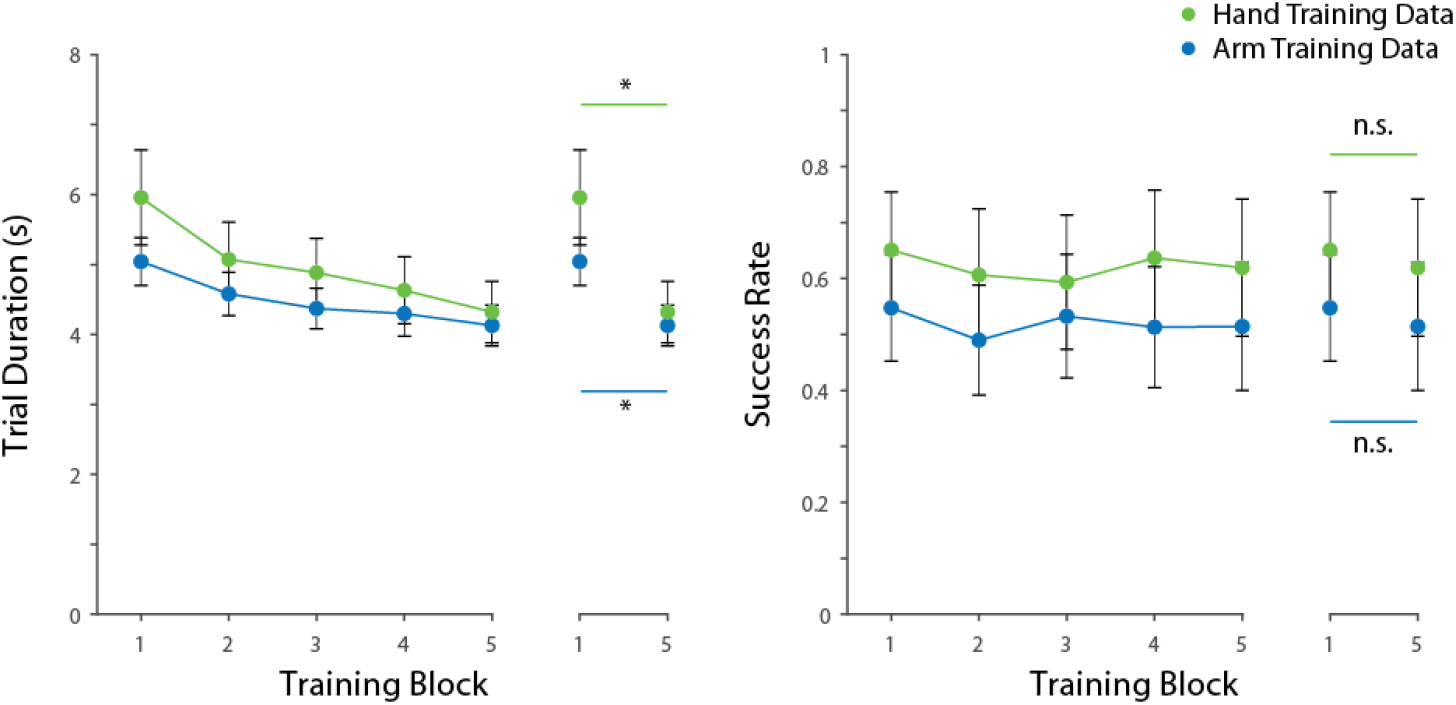
Training Data. A) Mean trial durations for the hand and arm training groups. Inset on right shows comparison between the first and final blocks of training for each group. B) Error rates for training blocks. Inset on right shows comparison between the first and final blocks of training for each group. Error bars present SEM. * Indicates p<0.01.

## Discussion

The present study examined transfer of motor skill within the same limb using an isometric force production task. We found that participants improved their ability to perform the task with both the trained and untrained effector. Specifically, participants who trained to perform the task using only their hand improved with both their hand and arm, and those who trained only with their arm improved with both their hand and arm. This effect was not due to learning during skill assessments, since a control group that did not train showed no changes in performance. We therefore conclude that the benefits of training on the task with one effector transferred to the untrained effector.

Notably, we found that training with either the hand or arm *alone* led to an improvement in skill when subsequently performing the task with *either* of these effectors. This non-effector-specific improvement is consistent with previous evidence for shared, generalized representations of motor skill (Schmidt 1975). In particular, previous research indicates a left hemisphere network of motor and premotor regions play a central role in motor skill learning (Reis et al. 2009; Hardwick et al. 2013b, 2015). Motor skill acquisition may therefore involve the development of a general control policy in left hemisphere motor regions that can be applied to perform tasks in an effector-independent manner. A further possibility is that training allowed participants to develop explicit strategies that could be applied when performing the task with either effector (Mazzoni and Krakauer 2006; Taylor et al. 2014). Our results provide no evidence of effector-specific training benefits on the skill task. Both training groups learned equally as shown by similar skill performance in the hand and arm assessments, despite training with only one of these effectors. This is of interest as the ability to control these effectors differs considerably. While we have fine control and are able to make precise movements with our hands, our arms are more suited to gross control, and the corticospinal representations of these effectors differ dramatically (Wassermann et al. 1992). Indeed, despite the similar magnitude of learning in both effectors, participants consistently had greater accuracy with their hand compared to their arm, indicating that controlling the cursor using the arm represented a more difficult task. As theories propose that more challenging training leads to greater improvements in performance (Guadagnoli and Lee 2004), it may have been expected that participants who trained using the more difficult arm task may have benefitted more from it. However, evidence from a recent study of skill-learning indicates no difference in performance between groups that trained on either an easy (slow) or difficult (fast) version of a skill acquisition task (Shmuelof et al. 2012). Thus, while our data look at the specific context of transfer, our results are consistent with recent work indicating that the difficulty of training does not have a significant impact on the final level of performance achieved.

Our results highlight the importance of considering skill acquisition as an improvement in a speed-accuracy trade-off, rather than improvements in the individual variables of speed and accuracy. During training, both the hand training and arm training groups improved their speed in performing the task, while maintaining a constant level of accuracy. However, during skill assessments, in which the speed at which they were required to perform the task was fixed, both groups performed with greater levels of accuracy than at baseline. This matches the pattern of results previously found in a study which examined learning with the arm version of this task in healthy individuals and stroke patients; improvements in speed during training can be translated to improved accuracy during skill assessments (Hardwick et al. 2017). This indicates that training led to an improvement in the ability to trade-off speed and accuracy, rather than a specific improvement in the individual variables of speed and or accuracy alone. Differences in self-selected training speeds also make it difficult to directly compare performance between groups during the training conditions. For example, the hand training group were slower, but more accurate, than the arm training group; therefore, from the training data alone, it would not be possible to determine whether one group performed better overall than the other during training. Using separate skill assessments that controlled the speed at which participants performed the task allowed us to provide a like-for-like comparison between these groups, again highlighting the importance of considering the relationship between speed and accuracy when examining performance (Fitts and Peterson 1964; Reis et al. 2009; Shmuelof et al. 2012; Hardwick et al. 2017, 2018).

The results of the present study contrast previous work on transfer of learning in visuomotor adaptation, which found learning generalized from the arm to the hand, but not from the hand to the arm (Putterman et al. 1969; Hay and Brouchon 1972; Krakauer et al. 2006). Krakauer et al., (2006) proposed that generalization depends on the context and prior history of action of the trained effector; all movements of the arm change the state of the hand, allowing shared learning, but not all movements of the hand affect the state of the arm, diminishing the likelihood of information learned with the hand to transfer. In comparison, the present study found transfer occurred when participants trained with either of these effectors. A primary difference between these studies is the nature of the tasks examined. As noted previously, adaptation involves modifying existing control policies, while motor skill learning involves developing new ones. Thus, while both skill acquisition and visuomotor adaptation examine ‘motor learning’, the processes and brain mechanisms underlying them differ considerably. Motor adaptation is largely attributable to cerebellum-dependent sensorimotor recalibration processes responsible for error detection/correction(Hardwick et al. 2013a), and is primarily governed by a cerebellar-prefrontal loop (Keisler and Shadmehr 2010; Liew et al. 2018). By contrast, skill learning recruits a broader network of functions and brain regions, including not only cerebellar-prefrontal error detection/correction mechanisms, but also sensorimotor-striatal reinforcement learning mechanisms (Hardwick et al. 2013b; Morris et al. 2016) and sensorimotor cortical plasticity mechanisms (Pascual-Leone et al. 1995; Mawase et al. 2017). Thus, compared to motor adaptation, skill learning recruits additional physiologically and functionally separate networks, including greater reliance on memory functions (Robertson 2007; Wollenweber et al. 2014; Shadmehr and Krakauer 2008). The more extensive network of brain regions and learning mechanisms used in the present skill task may explain these differences. A second key difference between these studies is that the present study did not involve movements; participants performed isometric contractions. As such, the relationship between movements of the effectors may have been of little consequence, as no movements were performed during the experiment.

## Conclusions

We examined intra-limb transfer in participants who learned to skillfully control the position of a cursor by producing isometric forces using either the hand or arm. Participants who trained with either effector improved their performance of the task with both their hand and arm. Participants who trained with their arm showed a trend for a larger improvement in their final skill with their arm compared to their hand, indicating some additional benefits may arise from training with effectors that have less initial fine control. These results indicate that training on a skill leads to the development of a shared control policy that can be generalized to different effectors. We propose this difference from the asymmetric transfer seen in visuomotor adaptation occurs due to the differences in the underlying learning processes (i.e. modification of existing actions vs establishing new ones). Finally, the ability to transfer the skill across effectors might be beneficial to develop training protocols for patients with motor impairment in specific effectors.

## Acknowledgements

PC and RH were supported by National Institutes of Health grant R01HD073147. RH was supported by a Marie-Sklodowoska Curie Individual Fellowship NEURO-AGE (702784).

Author Contributions
RMH, VAR, and PAC conceived the study. VAR collected the data. VAR and RMH analyzed the data. RMH drafted the manuscript. RMH, VAR and PAC finalized the manuscript.

